# i2dash: Creation of Flexible, Interactive and Web-based Dashboards for Visualization of Omics-pipeline Results

**DOI:** 10.1101/2020.07.06.189563

**Authors:** Arsenij Ustjanzew, Jens Preussner, Mette Bentsen, Carsten Kuenne, Mario Looso

## Abstract

Data visualization and interactive data exploration are important aspects of illustrating complex concepts and results from analyses of omics data. A suitable visualization has to be intuitive and accessible. Web-based dashboards have become popular tools for the arrangement, consolidation and display of such visualizations. However, the combination of automated data processing pipelines handling omics data and dynamically generated, interactive dashboards is poorly solved. Here, we present i2dash, an R package intended to encapsulate functionality for programmatic creation of customized dashboards. It supports interactive and responsive (linked) visualizations across a set of predefined graphical layouts. i2dash addresses the needs of data analysts for a tool that is compatible and attachable to any R-based analysis pipeline, thereby fostering the separation of data visualization on one hand and data analysis tasks on the other hand. In addition, the generic design of i2dash enables data analysts to generate modular extensions for specific needs. As a proof of principle, we provide an extension of i2dash optimized for single-cell RNA-sequencing analysis, supporting the creation of dashboards for the visualization needs of single-cell sequencing experiments. Equipped with these features, i2dash is suitable for extensive use in large scale sequencing/bioinformatics facilities. Along this line, we provide i2dash as a containerized solution, enabling a straightforward large-scale deployment and sharing of dashboards using cloud services.

i2dash is freely available via the R package archive CRAN.

## Introduction

Interactive data visualization is of vital importance when dealing with complex or extensive data as regularly produced in the fields of physics, geography or life sciences. In this context, recently published software packages providing interactive data visualizations in a web browser, such as Shiny (https://shiny.rstudio.com/) or Plotly [1], have become popular. Many newly developed software tools utilizing these “meta-level” packages were introduced, intended for the examination of specific aspects of data. Focusing on the R programming environment in the field of life sciences, examples of such software include tools that create static HTML reports for quality control and data set comparison, such as scRNABatchQC [2], or interactive R/Shiny based applications that allow data exploration and visualization, such as Cerebro [3], iSEE [4], and pcaExplorer [5]. Other packages, like WILSON [6], provide generic Shiny and Plotly based modules and data handling functionality, enabling the construction of customized web-apps. However, all these tools are restricted to a distinct data object in a certain format, e.g an R SingleCellExperiment object [7] in case of iSEE. Although such format dependencies ensure high performance of the visualization tools, they limit computational biologists to predefined plots, types of visualization, and fixed layouts. A generic software package able to be integrated in any kind of analysis pipeline, enabling web-based, custom defined layouts for data presentation and exploration, is critically missing. Here we introduce i2dash, a tool that provides functionality for data analysts working with any kind of omics data using complex pipelines. i2dash enables the programmatic creation of interactive dashboards from within such pipelines. In contrast to other applications such as scRNABatchQC and Cerebro, there is no need to define the final dashboard layout at the beginning of the pipeline. Instead, an iterative and dynamic process of dashboard creation is started, that facilitates a focus on story-telling suitable for diverse pipeline outcomes during runtime. Using high-level functions, i2dash liberates the data analyst from the struggle with low-level code or time consuming spatial arrangement of data visualizations.

## Results

### i2dash integrates arbitrary R workflows and communication on data

i2dash primarily acts at the interface of non-computational scientists (NCS) and computational data analysts (CDA) (**Figure 1**). It provides functionality for data analysts working with any kind of Omics data using complex pipelines. i2dash enables the programmatic creation of interactive dashboards from within such pipelines. In contrast to other applications such as scRNABatchQC and Cerebro, there is no need to define the final dashboard layout at the beginning of the pipeline. Instead, an iterative and dynamic process of dashboard creation is started, that facilitates a focus on story-telling suitable for diverse pipeline outcomes during runtime. Using high-level functions, i2dash liberates the data analyst from the struggle with low-level code or time consuming spatial arrangement of data visualizations.

**Figure 1.**
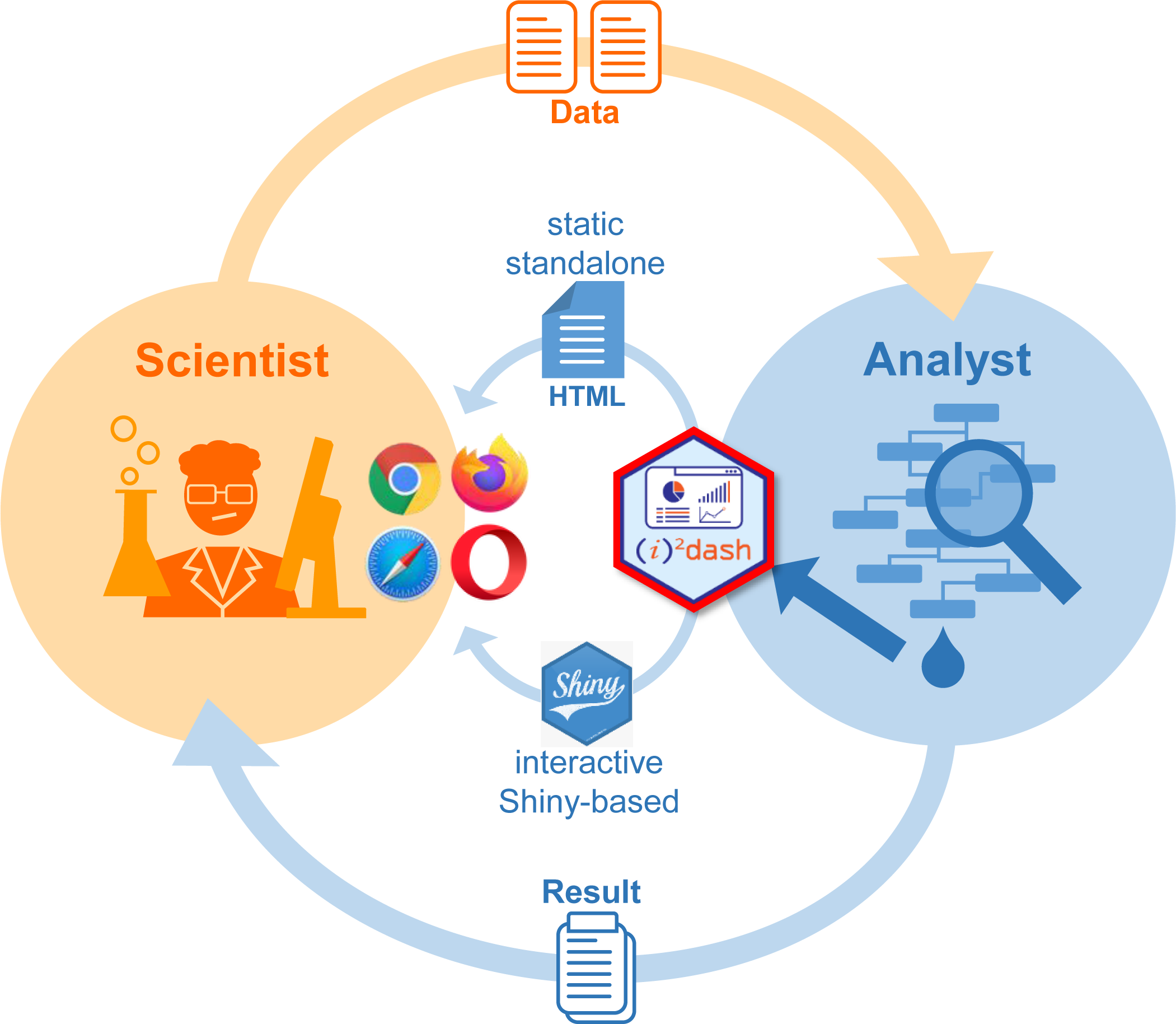
i2dash at the interface of scientist and analyst. Scientists and data analysts are typically connected by a process of data generation, data analysis, provisioning of results, and an iterative process of analysis refinement (orange and blue, outer arrows). i2dash (red hexagon) supports this process. It interacts with any kind of R based analysis pipeline, and provides functionality to generate both a static HTML output or an interactive R Shiny app. In order to access the data interpretation provided by the analyst, the scientist depends on a web-browser of choice (inner circle).

It is entirely written in the R programming language and relies on the widely used R packages flexdashboard (https://CRAN.R-project.org/package=flexdashboard), knitr [8] and RMmarkdown [9]. i2dash introduces a new class based on R’s S4 object system named i2dashboard, which by design provides the main functionality of the package. Besides global properties such as the dashboard title, author and theme, an instance of the i2dashboard class also stores individual dashboard pages, sidebars, navigation bar items, and document wide colormaps, as well as all components that make up the content of individual pages. i2dash has been tested on macOS, Linux and Windows. The latest stable version can be installed from the Comprehensive R Archive Network (CRAN, https://CRAN.R-project.org/package=i2dash) and the development version can be found at https://github.com/loosolab/i2dash. Moreover, i2dash is available as a Docker container in order to facilitate automated deployments (see below, https://gitlab.gwdg.de/loosolab/container/i2dash.deployment) and dashboard development in isolated environments, thus reducing the burden of software version management.

i2dash is designed to provide the generated dashboards both as html websites and interactive Shiny apps. Since the latter requires a local server or an advanced IT infrastructure for even larger deployments (e.g. in cloud environments), we provide a cloud compatible docker container to assist data analysts with dashboard distribution. Aside from usability, i2dash is designed to work with custom extensions for specific data types in order to generate adapted visualization venues in research and industry. As a proof of principle, we developed a single-cell RNA-sequencing (scRNA-seq) extension, enabling the generation of dashboards optimized for a typical single-cell (SC) data analysis workflow with very few function calls, providing interlinked visualizations among others (see use case below). From an end-user’s point of view (**Figure 1**), i2dash provides simplified access to high dimensional data interpretations via a web-browser. Hence, i2dash is the first choice to supersede static reports typically generated by many analysis pipelines by state of the art responsive, interactive and flexible dashboards.

### i2dash workflow

A typical workflow for the creation of a dashboard with i2dash is a flexible process, suitable for integration into existing analysis pipelines, that generates results iteratively (**Figure 2a**). Once an instance of a dashboard has been initialized, CDAs can add pages with various layouts (**Figure 2b**), enabling different content presentation strategies (see **Table 1**). Individual components such as texts, images, tables, and interactive widgets can be added to pages as defined by the inherent layout (see **Table 1**). Noteworthy, it is possible to link components into responsive views, such that data points visualized and selected in one component can trigger a change in the other. Dashboard creation concludes with the assembly of an RMarkdown document from the i2dashboard instance, which can be directly rendered into a static HTML file (e.g. for archiving or sharing with colleagues) or into an interactive Shiny app running on the local computer for web-based exploration (**Figure 2c**).

**Table 1.** Available dashboard layouts and features.

| Layout | Features | Useful for |
| --- | --- | --- |
| Storyboard | unlimited capacity | presenting a sequence of data visualizations and related commentary |
| Tabset<br>Column | unlimited capacity | presenting a large number of components without requiring the user to scroll |
| Focal left | up to three components<br>linkable layout | highlighting a single component |
| 2x2 grid | up to four components<br>linkable layout | cross-linked components / interactivity |
| Sidebar | globally or locally | providing static content and UI elements to control interactive components |

**Figure 2.**
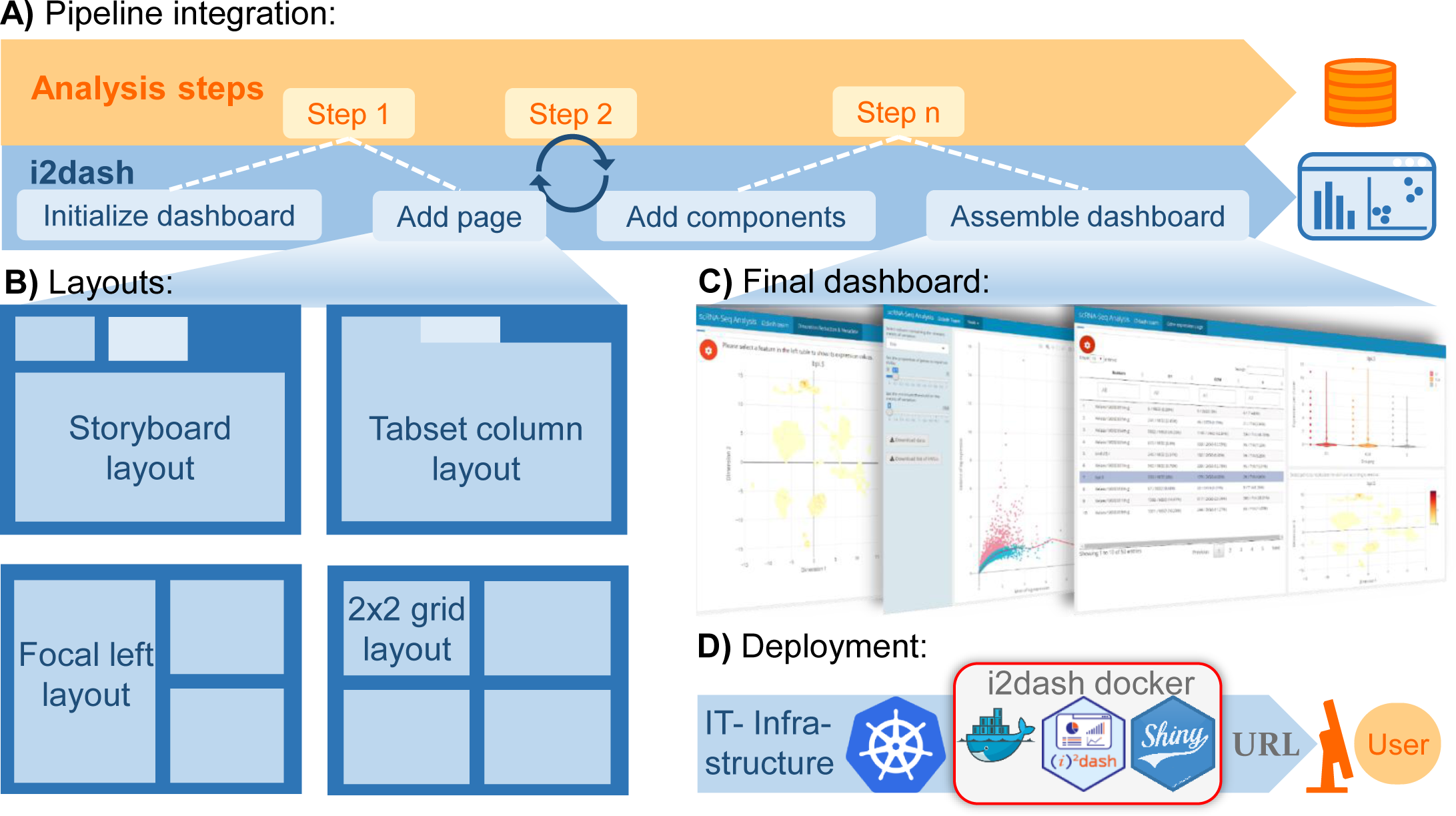
i2dash mechanism of action. **A.** A dashboard is initialized (blue arrow left) at the beginning of a data analysis pipeline (orange arrow, left). During the pipeline run, new content, results, or data visualizations are iteratively added to the dashboard’s pages. The final dashboard is rendered into a static or interactive document. **B**. Pages added to the dashboard can be arranged in a flexible manner from a selection of predefined layouts. **C**. Examples of programmatically created dashboards. **D**. The i2dash docker container, providing all dependencies for interactive, Shiny based apps, can be used to deploy individual data interpretations on a cloud infrastructure, such as Kubernetes, as a micro service.

### Add content to an i2dash dashboard

In order to achieve maximum customizability, we provided i2dash with a versatile mechanism to add content. Its generic functions can dispatch arguments based on their individual class(es). For example, the same generic function might add an image or an HTML widget to the dashboard, depending on whether it gets passed a file path or an object of a supported HTML widget class. This feature allows i2dash to handle objects of more than 50 different classes (for a complete list, see gallery.htmlwidgets.org). Moreover, i2dash implements the ability to pass functions as an argument to generic functions that are only expected to return plain RMarkdown code upon execution. Those functions allow the injection of arbitrary and user-defined content into the dashboard, constituting the most flexible mechanism conceivable.

### Cloud deployment of dashboards

Serving a dashboard on a local computer is simple and satisfies basic user needs. However, integration of dashboards into professional service facilities requires more advanced infrastructure features. Accordingly, we provide a docker container for deployment of interactive dashboards on local servers or even cloud infrastructure platforms such as Kubernetes (**Figure 2d**). In a typical use case, a service facility provides a running instance of the interactive dashboard to collaborators for interactive exploration of their data analysis. Following the FAIR data principles, access to the interactive dashboard can additionally be granted to the public audience, e.g. as an accompanying supplementary resource upon publication (as an example, see http://mpibn-mampok.134.176.27.161.xip.io/use-case-1/i2dash/).

### Use Case: single-cell RNA-seq data exploration with i2dash.scrnaseq

As indicated above, i2dash can be extended with functions for custom visualizations and tools. As a proof of concept, we developed an extension of i2dash for the exploration of results from scRNA-seq, called i2dash.scrnaseq (https://gitlab.gwdg.de/loosolab/software/i2dash.scrnaseq). It is intended to provide data visualizations and tools, assisting typical steps performed during single□cell RNA□seq analysis. We analyzed a previously published dataset ([10], GSE127465) of human and mouse lung cancers with scater [11], following the current best practices in scRNA-seq analysis [12] (**Figure 3a**), and populated an interactive dashboard using i2dash.scrnaseq.

**Figure 3.**
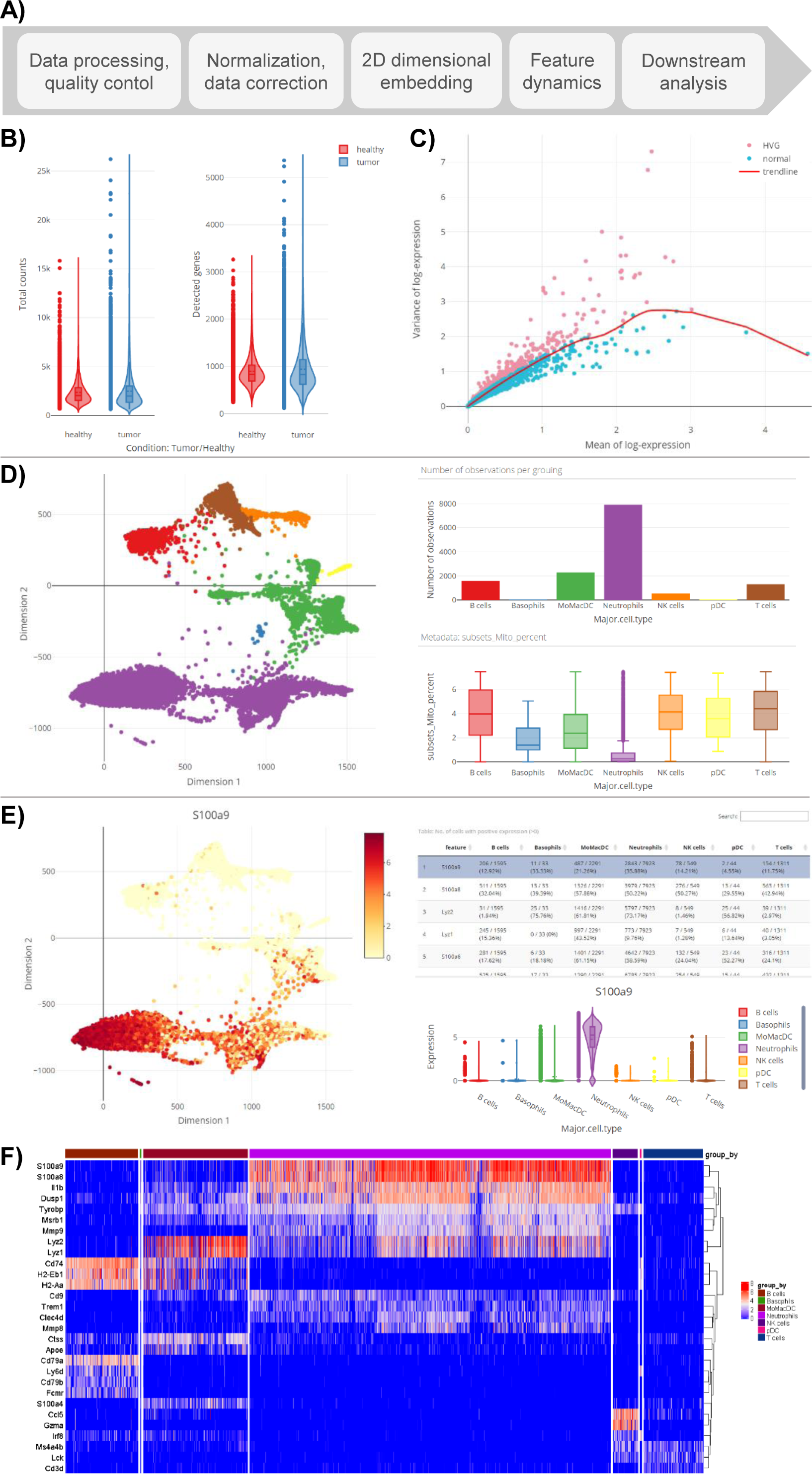
Visualization features of the i2dash.scrnaseq extension. **A**. High level workflow for SC analysis **B**. Violin plots from the Plotly package showing per-cell quality criteria, such as the number of counts and the number of detected genes. Plots are stratified by the categorical indicator “Condition”. **C**. Visualization of expression mean and variance of features (mean-variance plot). Highly variable features are highlighted and have been selected for downstream analysis. **D**. Exploration of cell metadata using linked visualizations. On the left, individual samples are shown in a 2D embedding, colored by their annotated cell type. Upon sample selection, the number of observations per cell type (top right) and the percentage of mitochondrial content across cell types (bottom right) are updated respectively. **E**. Exploration of feature metadata using linked visualizations. On the left, the feature expression of individual samples is shown in a 2D embedding. A table (top right) shows features of interest (e.g. from differential expression) and can update the coloring of the 2D embedding, as well as violin plots showing feature expression across cell types (bottom right). **F**. A heatmap is utilized to illustrate the top 10 marker genes by the annotated cell type.

Starting with per-cell quality control of the pipeline, i2dash.scrnaseq provides boilerplate functions for boxplots, barplots, or violinplots (e.g. to visualize the number of detected genes per cell) (**Figure 3b**). Optionally, those plots can be stratified by categorical variables (e.g. sequencing batch), to investigate whether data correction is needed (**Figure 3b**). Per-cell quality data can additionally be aggregated across levels of categorical data (e.g. to obtain the mean number of genes detected by sequencing batch) using any function to summarize numerical data (e.g. mean, median, sum, etc.). Feature selection of e.g. highly variable genes can be visualized using a mean-variance plot on the “Feature selection page” (**Figure 3c**).

Downstream analyses like cell clustering are often based on results from dimension reduction methods (e.g. t-SNE or UMAP). Consequently, high-level functions from i2dash.scrnaseq center around 2D embeddings (**Figure 3d, e**), allowing the exploration of metadata either on sample-level (e.g. the number of cells per inferred cell type, **Figure 3d**) or feature level (e.g. the expression of a marker gene in cell clusters, **Figure 3e**). The “Feature expression page” can additionally display results from differential expression analysis in a table that is linked to the 2D embedding and will interactively update coloring upon feature selection. To complete the portfolio, i2dash.scrnaseq provides functionality to display feature dynamics in an interactive heatmap (**Figure 3f**).

Besides high-level functions for the visualization and interactive exploration of results from scRNA-seq analysis, i2dash.scrnaseq provides helpful tools to simplify otherwise cumbersome tasks: The “Feature grid page” can be used to interactively create publication-ready graphics that shows the 2D embedding, color-coded by feature expression, for an arbitrary number of features in a grid layout (**Figure 4a**). In the context of dimension reduction methods, the “DimRed comparison page” simplifies the parameter search by comparing many different dimensionality reductions across different parameters (**Figure 4b**).

**Figure 4.**
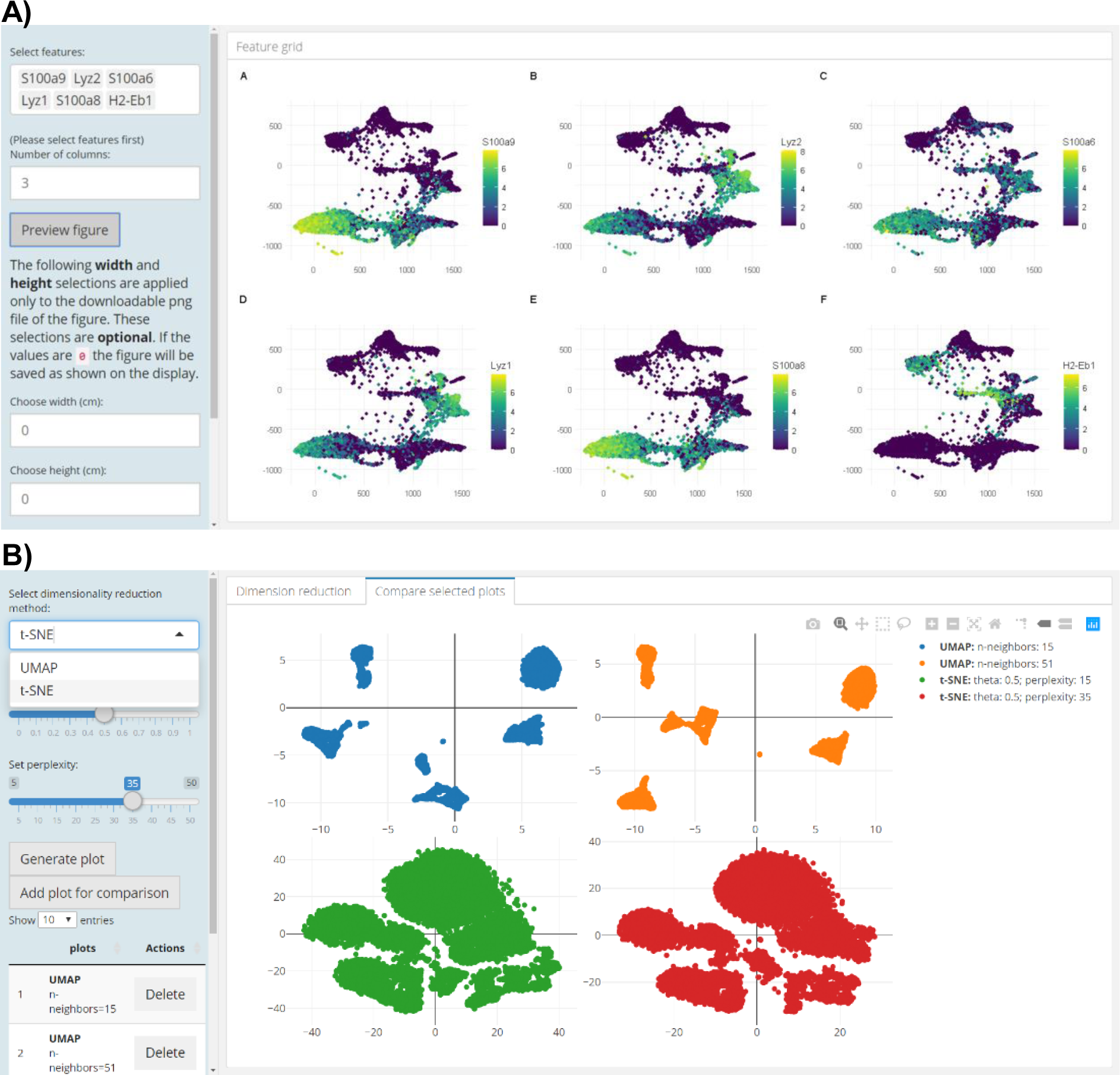
Tool features of the i2dash.scrnaseq extension. **A**. The “feature-grid-page” provides functionality to illustrate 2D embedded samples colored by expression of arbitrary features. The number of grid columns and figure sizes can be customized and the resulting figure, as well as the visualized data are provided for download. **B**. The “dimred-comparison-page” allows comparison of multiple 2D embeddings obtained by different runs of UMAP or tSNE. Parameters can easily be defined using sliders and results can be added individually to the comparison overview.

### Documentation and tutorials

We provide comprehensive documentation of i2dash and the S4 class i2dashboard online (https://loosolab.github.io/i2dash) and in the respective package vignettes. Documentation for each function of i2dash and its extension i2dash.scrnaseq is also provided using the man page system of R. Notably, package vignettes address users at different experience levels, addressing both beginners, intermediates, and experts in R. In addition, we provide comprehensive tutorials on how to use available functions from i2dash to extend functionality in order to allow R experts to fit dashboards to the field and data they work in.

### Benchmarking and feature comparison

The R package i2dash is a generic tool for the generation of flexible dashboards. Therefore, it cannot be directly compared to specialized applications as enumerated above, dealing e.g. only with single-cell specific aspects such as iSEE does. However, by including the SC specific extension i2dash.scrnaseq, a comprehensive comparison becomes feasible, emphasizing the generic approach of our framework. A detailed side-by-side comparison of web-based visualization tools from the perspective of implemented features is shown in **Table 2**. Briefly, all compared tools are R-based applications, able to be deployed on a multitude of operating systems. In terms of the general customization, i2dash offers a wider range of layouts and ways to organize content. While iSEE does allow a customized organization of visualization panels, i2dash extends this feature by content organization via pages, menus, and sidebars. With regard to the content type, the other programs are limited to SC specific data types, provided as R objects. Only i2dash supports a variety of content types, including but not limited to images, text, and HTML widgets. Interactive and linked visualizations embedded in a Shiny application are provided by almost all programs. However, i2dash offers the unique possibility to link interactive visualizations in a customizable way, a feature of high importance when specific aspects of data should be prioritized. Regarding the SC related features, i2dash is capable of providing all key aspects of a common scRNA-seq analysis. In summary, we conclude that i2dash and i2dash.scrnaseq at least substitute other specialized applications in the field of SC analysis, and define a new standard with respect to applicability in the context of customized, interactive, and web-based dashboard development.

**Table 2.**
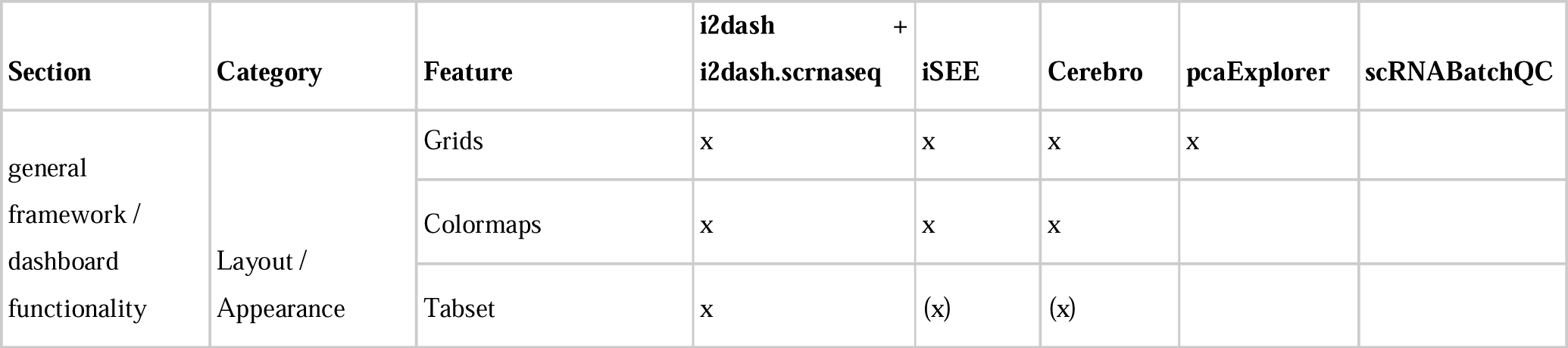
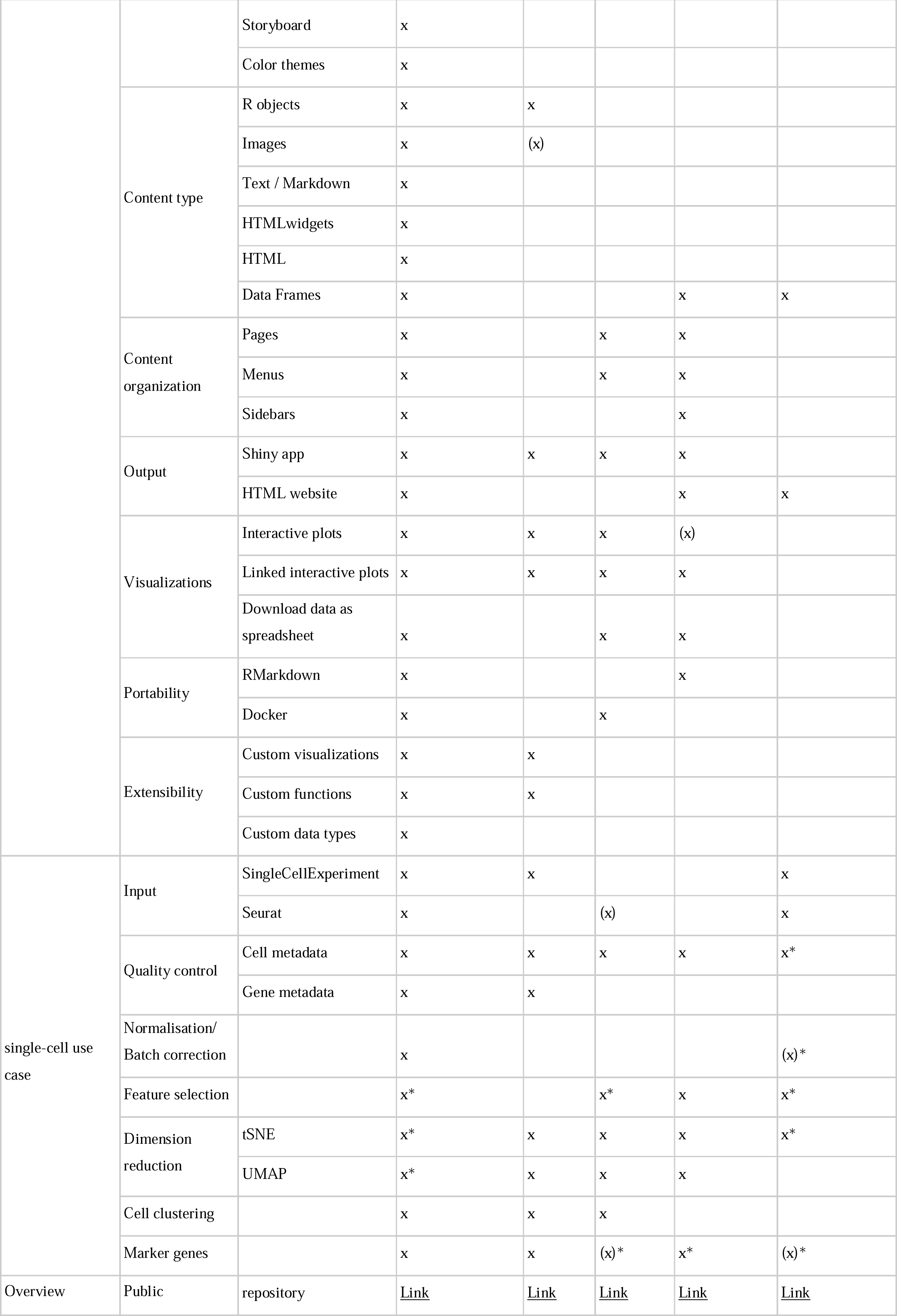
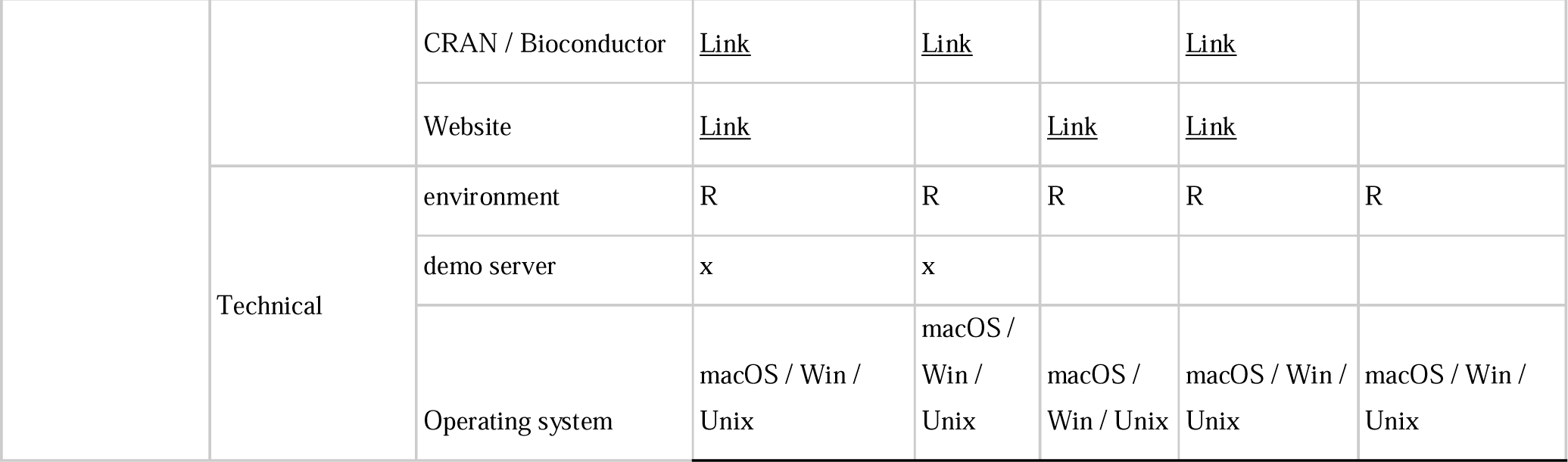
Benchmarking and feature comparison between web-based applications focusing on single-cell RNA-seq. *Notes*: “x” = implemented feature/functionality; “(x)” = limited or in similar form; “*” = calculated during the usage of the application.

### Conclusions

i2dash is an extensible R package for the creation of flexible dashboards by iterative addition of interactive data visualizations. i2dash integrates seamlessly into existing data analysis pipelines, supports responsive (linked) components, and the resulting dashboards can easily be shared or deployed onto cloud infrastructure. We extended i2dash for data from single-cell RNA-seq experiments, enabling sample quality control, exploration of metadata and feature expression, visualization of feature dynamics, and the creation of publication-ready graphics.

## Materials and methods

### Availability and requirements

**Project name:** i2dash, i2dash.scrnaseq

**Project home page:** https://gitlab.gwdg.de/loosolab/software/i2dash

**Project documentation:** rendered at https://loosolab.github.io/i2dash

**Operating system(s):** Linux, macOS, Windows

**Programming language:** R

**Other requirements:** R 3.5 or higher, Bioconductor 3.12 or higher, Pandoc

**License:** MIT

**Any restrictions to use by non-academics:** none.

**Container image:** https://gitlab.gwdg.de/loosolab/container/i2dash.deployment

### Demos

http://mpibn-mampok.134.176.27.161.xip.io/use-case-1/i2dash/

http://mpibn-mampok.134.176.27.161.xip.io/use-case-2/i2dash/

http://mpibn-mampok.134.176.27.161.xip.io/use-case-3/i2dash/

## Authors’ contributions

JP and ML conceived the design of the tool. AU and JP performed programming and bio-informatics analysis of use cases. MB and CK contributed to design, discussions and gave advice. All authors wrote the manuscript.

## Acknowledgements

This work was funded by the Deutsches Zentrum für Herz-und Kreislaufforschung (DZHK, Rhein-Main Site) and the Cardiopulmonary Institute (CPI).

## Supplementary Material

**Table S1.**
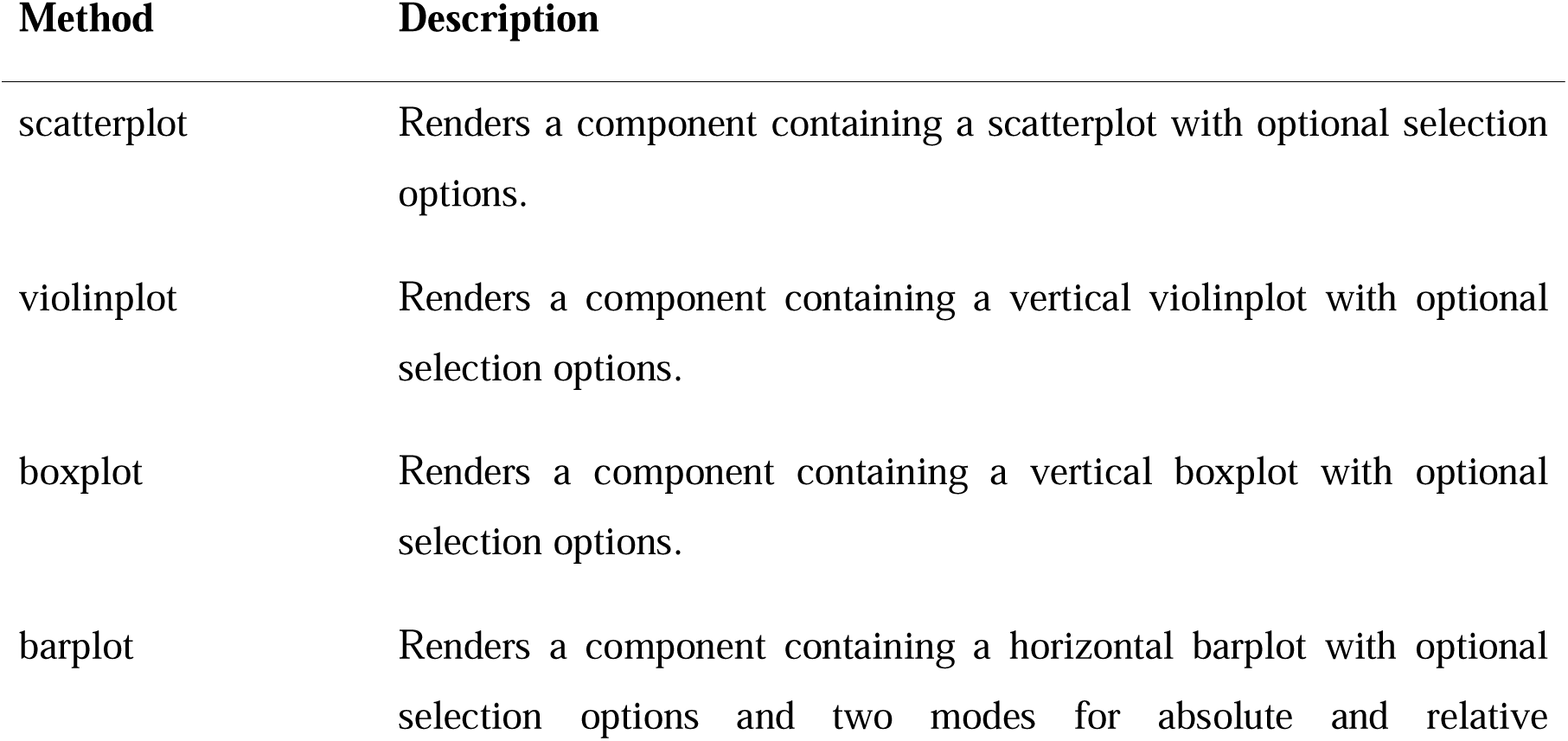
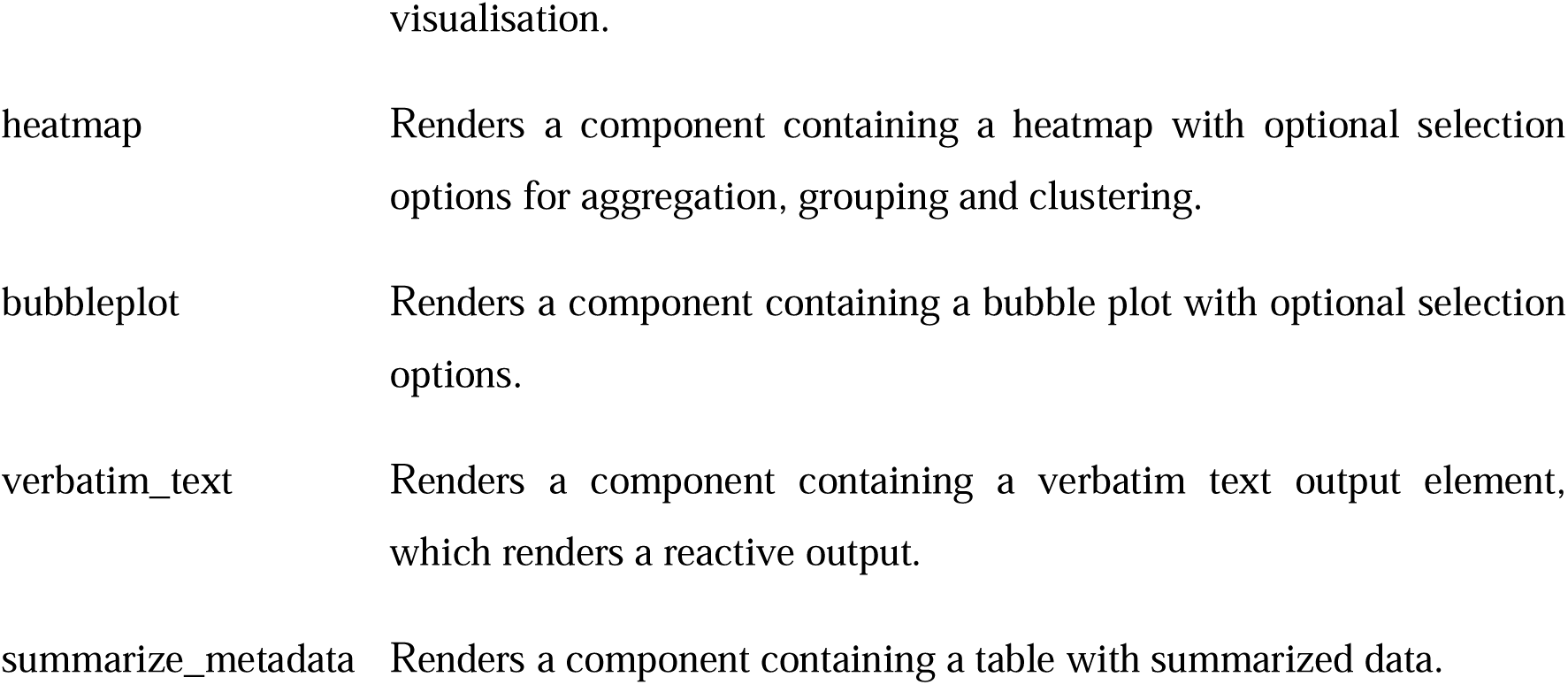
Component-methods of the package i2dash.scrnaseq.

**Table S2.** Pre-defined page-methods of the package i2dash.scrnaseq.

| Method | Description |
| --- | --- |
| add_dimred_feature_page | View a dimension reduction side-by-side with feature metadata. |
| add_feature_expression_page | Explore feature expression with dimension reductions, a violin plot and a table grouped by sample metadata. |
| add_dimred_sample_page | Characterize and visualize dimension reductions and sample groupings/ metadata. |
| add_feature_grid_page | Create expression visualization for multiple selected features on a regular grid. |
| add_dimred_comparison_page | Explore the effects of the parameters "theta" and "perplexity" of a t-SNE or "n_neighbors" of a UMAP embedding. |
| add_feature_selection_page | Quantify per-gene variation and explore the threshold on the metric of variation to get the desired set of highly variable features. |

